# Disentangling the reproductive and metabolic transcriptional responses to diet in *Drosophila melanogaster*

**DOI:** 10.1101/2025.10.16.682881

**Authors:** M. Florencia Camus, Avishikta Chakraborty, Max Reuter

## Abstract

Nutrition significantly influences various life-history traits in organisms, impacting decisions about growth, reproduction, and longevity. Previous studies in *Drosophila* have demonstrated that diet affects gene transcription, with many genes exhibiting altered expression between protein- and carbohydrate-rich diets. Some responses, such as the upregulation of oogenesis-related genes in protein-rich conditions, are physiologically intuitive, yet it remains challenging to distinguish between metabolic adaptations to different diets and regulation pertaining to reproductive investment in response to nutrient availability. In this study, we explore the transcriptomic responses of virgin and mated flies to changes in nutritional environments. We confirm previous phenotypic findings by observing responses to dietary preferences and nutritional needs. Our results indicate that both nutritional conditions and mating status lead to significant changes in gene expression in females. By comparing responses between virgin and mated flies, we differentiate between basal dietary responses and reproductive adaptations, with the latter involving eight times as many genes as the former. Notably, we identify GATA family transcription factors and the heat-shock factor (*Hsf*) as crucial regulators of diet-dependent reproductive genes. These findings enhance our understanding of the complex interactions between nutrition and reproductive strategies in *Drosophila*.

## Introduction

Nutrition plays a central role in life-histories across the animal kingdom, shaping traits such as growth, development, the onset of reproductive maturity, investment in gamete production, and survival. It does so by providing not only building blocks and energetic fuel for organisms, but also environmental cues that are used to make active decisions about how to modulate life histories to maximise expected lifetime fitness. One of the best-studied traits that is heavily dependent on nutrition is reproduction, where the quality and quantity of resources affect investment in energetically costly offspring. In sexual species, females in particular allocate significant resources to the production of a small number of large-sized gametes and accordingly nutrition has a large impact on female fecundity [1, 2]. For example, *Drosophila* females housed in higher protein environments produce up to four times as many eggs as females in carbohydrate-rich (and more protein-poor) environments [3].

Recent work in fruit flies assessing the transcriptional responses to diet suggests that diet-dependent variation in female fecundity is at least in part due to active regulation of reproductive investment rather than nutrient limitation [4]. Thus, gene expression changes observed between an optimal protein-rich and a suboptimal carbohydrate-rich diet mirror those observed in response to dietary restriction; similar to limiting dietary input, providing a diet with a sub-optimal macronutrient balance triggers an evolutionarily conserved life history response, where reproduction is actively down-regulated. This life history response relies heavily on the insulin-signalling and especially the TOR pathway [4–6], via members of the GATA transcription factor family [4, 7]. The presence of these responses makes it difficult to interpret diet-dependent physiological and transcriptional changes, and discriminate which transcriptional responses are to the diet itself, and which are a consequence of the altered reproductive investment.

One way to disentangle these effects on the female transcriptome, is to combine diet treatments with manipulation of a female’s mating status. In flies, like in most other organisms, a female’s first mating triggers the transition from a non-reproductive to a reproductive state [8–10]. This also brings about permanent physiological changes to be able to meet reproductive demands [11, 12]. For example, studies have documented a drastic shift in nutritional preference before and after mating; while virgin females prefer to feed on carbohydrate-rich foods similar to males, mated females not only consume more food, but switch to preferring diets higher in protein content in order to be able to meet the nutritional demands of egg production [13, 14]. Interestingly, this shift occurs even in genetically sterilized females that do not possess a germline and are unable to produce eggs [15]. In large parts, these changes are reliant on male-derived sex peptide, which is delivered as part of accessory gland fluids during mating [16–18]. Several transcriptomic studies have demonstrated that, as expected, the phenotypic changes after mating are associated with widespread responses at the gene expression level [15, 19–21]. But again, it is not clear to which degree these changes are a general, hard-wired response to mating and the onset of reproduction or diet-dependent, and if the latter, how expression changes triggered by mating are aligned with responses to diet.

Here, we report experiments that manipulate both diet and mating status in *Drosophila melanogaster* to dissect the complex relationship between nutritional and reproductive responses. Examining gene expression of virgin and mated flies fed either protein- or carbohydrate-rich synthetic diets, we identify and characterise expression changes that represent physiological adaptations to diet (shared by all flies), those that are generic responses to mating (invariant across diets) and those that modulated by both mating status and diet composition. We further compare these classes functionally and in terms of their regulation.

We find several hundred genes respond to mating and nutrition. A large proportion of genes show diet-dependent expression changes that are similar between virgin and mated flies. These genes include a set of core metabolic genes involved in physiological diet responses that are independent of mating status. But we also find evidence for a larger set of genes where expression differences tend to be more pronounced in the mated cohort. Concordant with previous work, we show that the diet response is mainly regulated by GATA transcription factors. However, we also identify *Hsf* (heat shock factor) as an important transcription factor modulating the upregulation of diet-specific reproductive response. In line with a role of *Hsf*, additional experiments reveal effects of mating status and diet on resistance to heat shock, adding support to an emerging role of heat shock factors in reproduction.

## Materials and Methods

### Fly Stocks and Maintenance

We used *D. melanogaster* from the laboratory population LH^M^. This population has been maintained as a large outbred population for over 400 non-overlapping generations [22, 23]. It is kept on a strict 14-day regime, with larval densities kept constant (∼150–200 eggs per vial containing ca. 7ml of media) and a fixed adult population size (56 vials of 16 male and 16 females) at each generation. Flies were reared at 25^°^C, under a 12h:12h light:dark regime, on a cornmeal-molasses-yeast-agar food medium (see **Supplementary Methods**). For all experiments involving capillary feeding, we used high relative humidity (>80%) to minimise evaporation.

### Synthetic Diet

For experiments, we used a modified liquid version of the holidic diet described in Piper et al. [24]. This food is prepared entirely from synthetic components to enable precise control over nutritional value (see **Supplementary Methods**). Two artificial liquid diets were made which differed in the ratio of protein (incorporated as individual amino acids) and carbohydrate (sucrose); with all other nutritional components provided in fixed concentrations. The nutrient ratios [P:C] used were 2:1 and 1:4, with the final concentration of each diet being 32.5g/L. These ratios were identified in previous work [4] as maximising the reproductive fitness of female and male LH^M^ flies, respectively. We note that P:C ratios may not be directly comparable to standard fly media, as nutrients in synthetic diets appear to be more readily accessible [24].

### Phenotypic data collection

*Diet preference:* Flies from each sex were collected as virgins using CO^2^ anaesthesia. Triplets of virgins were placed in individual vials containing molasses-yeast-agar culture medium (see Supplementary Methods for full recipe) with no added live yeast. We used triplets of flies as a means to balance between, on the one hand, replication across vials and, on the other hand, between-vial variance and workload [25]. Twenty vials of triplets were collected for each sex and mating status. All flies were aged for 2 days for them to become sexually mature. Subsequently, females in the “mated” treatment were provided with three virgin males (collected at the same time as females) as mating partners. These hextets of flies were left for 5 hours to mate, before being re-split by sex and transferred to new vials containing an 0.8% agar-water mixture. Virgin flies were not perturbed during this period and placed on the 0.8% agar medium at the same time as mated flies. The agar-water vials provide water for the flies but have no nutritional value.

In accordance with previous literature using this methodology [25, 26], flies were kept in agar-water vials overnight, then supplied with two 5µl microcapillary tubes (ringcaps©, Hirschmann) containing amino acid and carbohydrate solutions, respectively. Capillary tubes were replaced daily, and food consumption for each fly trio was recorded for a period of three days. As a control, the rate of evaporation for all diet treatments was measured in six vials that contained the two solution-bearing capillary tubes but no flies, placed alongside the experimental vials in the controlled temperature room. Their average evaporation per day was subtracted from the values of volume consumed in experimental vials to correct diet consumption for evaporation.

*Nutritional requirement:* The setup for nutritional requirement is almost identical to the diet choice experiment, except at the point of synthetic diet allocation. Instead of two capillary tubes (as per diet choice), flies were supplied with one microcapillary tube (ringcaps©, Hirschmann) containing one of the two allocated diets. These diets varied in their protein-to-carbohydrate ratios and captured the following nutritional rails (P:C): 2:1, 1:4. We had 20 vials with fly triplets per allocated diet and mating status. Capillary tubes were replaced daily, and food consumption for each fly trio was recorded for a total period of three days. Flies were exposed to diet treatments in a controlled temperature room (25°C), 12L:12D light cycle and high relative humidity >80%. The rate of evaporation for all diet treatments was measured by using five vials per diet that contained no flies, placed randomly in the constant temperature chamber. The average evaporation per day was used to correct diet consumption for evaporation.

*Female fitness:* Female adult fitness was measured as the number of eggs produced over a fixed period of time (18h). This performance proxy is expected to correlate closely with other fitness measures, such as the total number of offspring [27, 28]. Following the feeding period, trios of virgin or mated females were placed in new agar vials and presented with three males of the same age from the LH^M^ stock population. We note that this mating event is the second mating opportunity for the “mated fly” group, and this was done so that we could compare fecundity of recently inseminated females across both groups. Flies were left to mate/oviposit for 18 hours in vials containing *ad libitum* food corresponding to their diet treatment provided via capillary tubes. All flies were removed after this 18-hour mating window. Following removal of the flies, the total number of eggs laid were determined by taking pictures of the agar surface and counting eggs using the software *QuantiFly* [29].

*Male fitness:* Adult male fitness was quantified by counting the number of offspring produced in competitive mating trials, a metric previously validated in our lab as a reliable indicator of reproductive success, given that total adult progeny production in our population primarily depends on mating success [30]. Our method closely followed the experimental design outlined in [31], in which focal experimental males competed against standard competitor males for mating opportunities. After the previously described feeding period, a group consisting of three focal males (from both virgin and mated treatments), three virgin competitor males, and six virgin females was transferred to a fresh vial containing molasses-yeast-agar medium lacking live yeast, the primary food source for both sexes [32, 33]. Competitor males and females carried an LH^M^ genetic background and were homozygous for the recessive *bw*− eye-colour mutation. These competitor flies were raised under identical conditions and matched the age of the focal males. Following a 24-hour egg-laying period, adults were removed from the vials. The resulting eggs developed for 12 days, after which adult offspring were counted and classified based on eye colour to determine paternity: red-eyed offspring indicated paternity by the focal experimental males, whereas brown-eyed offspring were fathered by the competitor males.

*Thermal tolerance:* Fly thermal tolerance was measured only on female flies following established heat knockdown assays [34]. Following the feeding period in triplets, individual females were placed into 4ml water-tight glass vials, clipped onto a two-sided rack and immersed in a water bath with temperature set to 39°C. Thermal tolerance was measured as the time (minutes) it took an individual fly to stop locomotory activity (enter a coma-like state). The experiment was performed over two runs, with each run measuring heat knockdown time of 90 individual flies. Thus, a total of 180 flies were measured, corresponding to 15 fly triplets for each combination of diet and mating status.

*Statistical analysis:* To determine if dietary choices differed, we used a multivariate analysis of variance (MANOVA). The main model had protein and carbohydrate consumption as response variables, with mating status, sex, and their interaction as fixed effect. We performed subsequent univariate analysis of variance (ANOVA) to determine which nutrient(s) contributed to the overall multivariate effect. All analyses were performed using the *manova* function.

To examine whether the flies varied in the quantity they consumed of each diet, we used a linear model to investigate differences in dietary consumption. We modelled total food consumption as a response variable with diet treatment, sex, and their interaction as fixed effects. Female fitness was measured as total number of eggs produced within an 18-hour timeframe following a mating event. Male fitness was measured as the proportion of offspring sired from the focal male. For this we modelled the response as a binomial vector comprising the number of offspring sired by the focal male and the number sired by the competitor male and diet composition as a categorical fixed effect. Given data followed a normal distribution, we used a Gaussian linear model to analyse the data. Number of eggs was the response variable, with mating status, and diet plus their interaction as fixed factors.

Thermal tolerance data was analysed with a Gaussian mixed model, fitted with the *lmer* function in the *lme4* package [35]. The response variable was time (minutes) for flies to become unconscious, with mating status and diet (plus their interactions) as fixed effects. We included the nested term of “side of tank” within “run” of experiment as a random effect.

### Transcriptomic analysis

*Experimental Setup:* Flies from each sex were collected as virgins using CO^2^ anaesthesia. For the virgin treatment, groups of six virgin flies were collected and placed into vials containing culture medium (molasses-yeast-agar) with no added live yeast. For the mated treatment, three virgin females and three virgin males were placed in individual vials. For both treatments there was a total of six flies in the vials. Flies were left in these vials for a period of 36-hours in which the mated treatment had the chance to mate. Following this period, flies were split into triplets, and placed on 0.8% agar-water mixture. Agar-water vials provide water for the flies but have no nutritional value. Flies were kept in these vials overnight before being supplied with a microcapillary tube (ringcaps©, Hirschmann) containing one of the two allocated diets.

*Sample collection and RNA extraction:* For each sex, we generated 3 biological replicates for each diet treatment (virgins on protein diet, virgins on carbohydrate diet, mated on protein diet, mated on carbohydrate diet), a total of twelve samples. For each replicate sample, we pooled 4 triplets (a total of 12 flies) to ensure we collected sufficient RNA. Total RNA was extracted using the *Qiagen RNeasy Minikit* (Qiagen BV, Venlo, The Netherlands) according to manufacturer’s instructions. Quantity and quality of RNA was first inspected using a Nanodrop 2000 spectrophotometer (Wilmington, USA), and later verified using an Agilent Tapestation 2200 (Agilent, USA) at the UCL Genomics facility.

*Sequencing and read mapping:* Library construction and sequencing were performed at the UCL Institute of Child Health Genomics facility. Barcoded cDNA libraries were constructed using the *KAPA Hyper mRNA Library prep kit* (Roche, USA) and mixed at equal concentrations. This multiplexed sample was sequenced (43bp paired-end reads) on four flowcell lanes on an Illumina Nextgen 500 instrument to an average of 18M reads per sample.

Having verified that there was no bias towards particular libraries across the sequencing lanes using the Illumina Basespace online server, we merged reads from all four lanes. Adaptors and low-quality base pairs (below quality value of 3) were trimmed using trimmomatic v0.36 [36]. Trimmed reads from each sample were independently mapped to the *D. melanogaster* genome release 6.19 using HISAT2 [37]. Mapped reads were manipulated using *samtools* [38].

*Statistical analyses:* Exon-level read counts for each annotated gene were obtained using *htseq-count* [39], using release 6.19 annotations obtained from the ENSEMBL Biomart, and then summed across all exons within a single gene. Total read counts for each gene for the twelve samples were then used for differential gene expression analysis using the Bioconductor package *edgeR* [40], analysing each sex separately.

We first removed lowly expressed genes (read count < 2 across all samples). Read count data were then normalised across libraries and expression dispersion parameters calculated using the entire dataset. We tested for differential gene expression between our experimental groups using the negative binomial models implemented in *edgeR*. We fitted a full model where expression of each transcript was a function of mating status, diet and their interaction. The significance of each model term was tested using a specific contrast matrix. To obtain estimates of expression fold changes between the two diets for each mating status, we further fitted separate models for each mating status with diet as the sole term.

Gene ontogeny enrichment was performed using the Bioconductor package *clusterProfiler* [41]. We further compared our list of genes that responded to mating to previous work that has examined transcriptomic responses of changes in mating status [20]. For this, we used the R package *GeneOverlap* [42] uses contingency table tests to identify greater than expected overlap between gene lists. In order to assess whether genes that showed similar diet responses were regulated by common transcription factors we used the Bioconductor package *RcisTarget* [43], which tests for enrichment of cis-regulatory motifs in 5kb windows upstream of genes in a given sets. In all analyses, we used a statistical significance threshold of 5% False Discovery Rate (FDR) [44]. All analyses were performed in R version 3.3.2 [45].

## Results

### Phenotypic responses

Phenotypic data collected alongside our expression data recapitulated previously described patterns of diet preference and diet-dependent choice and fitness. Briefly, we observe sexual dimorphism for diet preference, diet consumption and diet-dependent fitness, with females both preferring and performing better on high-protein diets and males on high-carbohydrate diets (**Supplementary Results 1**). Preference assays showed that females, but not males, show a large shift in nutritional preference following mating. While in a choice assay virgin females consumed separately offered protein and carbohydrate diets in relatively small quantities and in proportions that resulted in a protein-to-carbohydrate ratio of 1:2.5 (not dissimilar to the composition chosen by males), mated females consumed a much larger total quantity of food and almost tripled their protein intake, resulting in a significantly altered protein-to-carbohydrate ratio of 1 : 0.92 (**Figure SR1.1A, Table SR1.1**). In no-choice assays, we found that mating has a significant effect on female total food consumption (**Figure SR.1B**, **Table SR1.2**), with mated females consuming more than virgin females. We also found that mated females consumed similar amounts when supplied with high-protein or high-carbohydrate carbohydrate diets, whereas virgin females consumed more on the high-protein diet (**Figure SR1.1B**, **Table SR1.2**). Reproductive fitness measurements showed sex-specific effects, whereby virgin females (freshly mated in our assay) were able to lay more eggs than mated females (who had already been previously reproductively in our assay) across both diets, but this fitness difference was not diet-dependant. Unlike in females, mating did not influence male fitness. Regardless of mating status however, females performed best on protein and males on carbohydrate (**Figure SR1.1, Table SR1.3**).

### Differential gene expression

After QC, our dataset included expression data for 12,880 genes for males and 9,862 genes for females. For males, we found 139 genes responding significantly in expression to changes in diet; no genes were differentially expressed in relation to mating or its interaction with diet (**Table 1**). Genes that were differentially expressed in relation to diet included a number of that are associated with male ejaculates, specifically accessory gland proteins (24A4, 53C14a, 53C14c, 53Ea, 62F, 63F, 76A), seminal fluid proteins (24Bb, 33A3, 53C14a, 60F, 87B) and serpins (28F, 38F, 77Bb), as well as *Seminal metalloprotease-1*, *Ejaculatory bulb protein* and *Ejaculatory bulb protein II* (see online archived data for a full list of genes [46]). All of these are down-regulated on protein-rich compared to carbohydrate-rich food, in line with the diet-specific reproductive investment observed in our phenotypic assays.

**Table 1:**
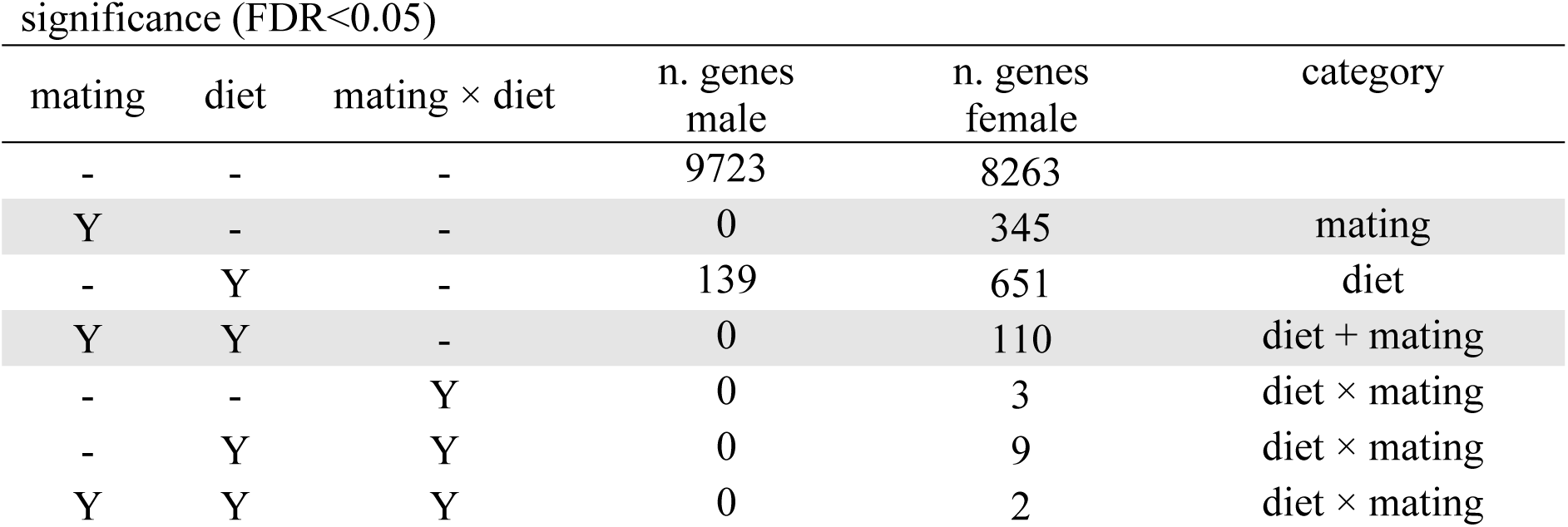
Numbers of genes that do or do not show significant differential expression in response to Mating, Diet, and their interaction in males and females. Based on the significance of the three predictor terms, genes were classified into four functional categories (last column).

| significance (FDR<0.05) |  |  |  |  |  |
| --- | --- | --- | --- | --- | --- |
| mating | diet | mating × diet | n. genes<br>male | n. genes<br>female | category |
| - | - | - | 9723 | 8263 |  |
| Y | - | - | 0 | 345 | mating |
| - | Y | - | 139 | 651 | diet |
| Y | Y | - | 0 | 110 | diet + mating |
| - | - | Y | 0 | 3 | diet × mating |
| - | Y | Y | 0 | 9 | diet × mating |
| Y | Y | Y | 0 | 2 | diet × mating |

Transcriptomic responses in females were considerably more complex (**Table 1**). We identified 651 genes that showed differential expression between the diet treatments (‘Diet’ set), 345 genes that showed differential expression between virgin and mated flies (‘Mating’ set), 110 genes that showed additive expression changes in response to both diet and mating (‘Diet+Mating’ set) and 14 genes that showed a significant diet-by-mating interaction (‘Diet×Mating’ set), some in combination with additive diet and/or mating effects.

Genes in the Diet set represent a core metabolic response to nutritional composition. Expression changes occur in both directions (up- and down-regulation when going from a carbohydrate to protein environment, **Figure 1A**) and carbohydrate-to-protein fold changes are positively correlated between virgin and mated flies (r = 0.826, p < 0.001). In line with these genes being part of a basal metabolic response to diet, female Diet genes significantly overlap with the genes that show a significant diet effect in males (overlap = 41 genes, p < 0.001). Furthermore, genes in the female Diet set overlap with genes that we previously found to show concordant diet responses between mated males and females (S category in [4]: overlap = 305 genes, p < 0.001, **Table 2**). But in addition, the Diet set also includes smaller numbers of genes that show female-specific responses to diet, and were previously shown to respond differently to diet between mated females and males (DxS category in [4]: overlap = 15 genes, p < 0.001; D+(D×S): overlap = 70 genes, p < 0.001, **Table 2**). GO analysis showed that Diet genes are significantly enriched for terms that relate to neuronal signalling and response to stimulus (**Figure 2**).

**Figure 1:**
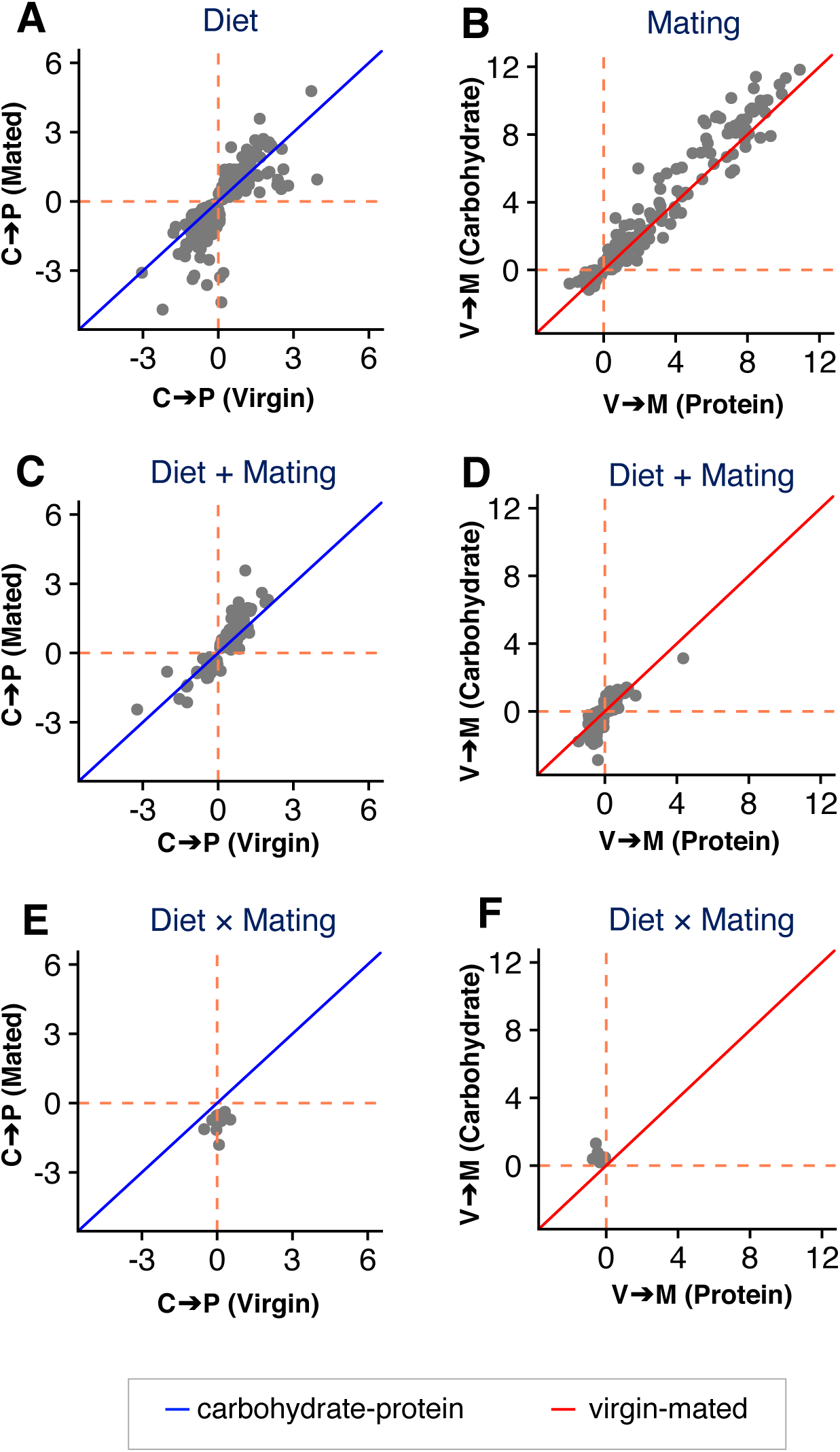
Correlations in log2-fold changes across treatments for genes classified as showing: only a diet effect (Diet) or only a mating effect (Mating) (first row of panels), an additive effect of both (Diet + Mating, second row) or a diet-by-mating interaction (Diet × Mating, third row). Fold changes in the left column are from the carbohydrate-rich to protein-rich diet and shown between virgins (x-axis) and mated females (y-axis), fold changes in the right column are from virgins to mated females and shown between the protein-rich diet (x-axis) and the carbohydrate-rich diet (y-axis). Orange dashed lines indicate zero-fold change for the two dimensions, blue and red diagonal lines designate equal fold change in the two conditions (slope = 1).

**Figure 2:**
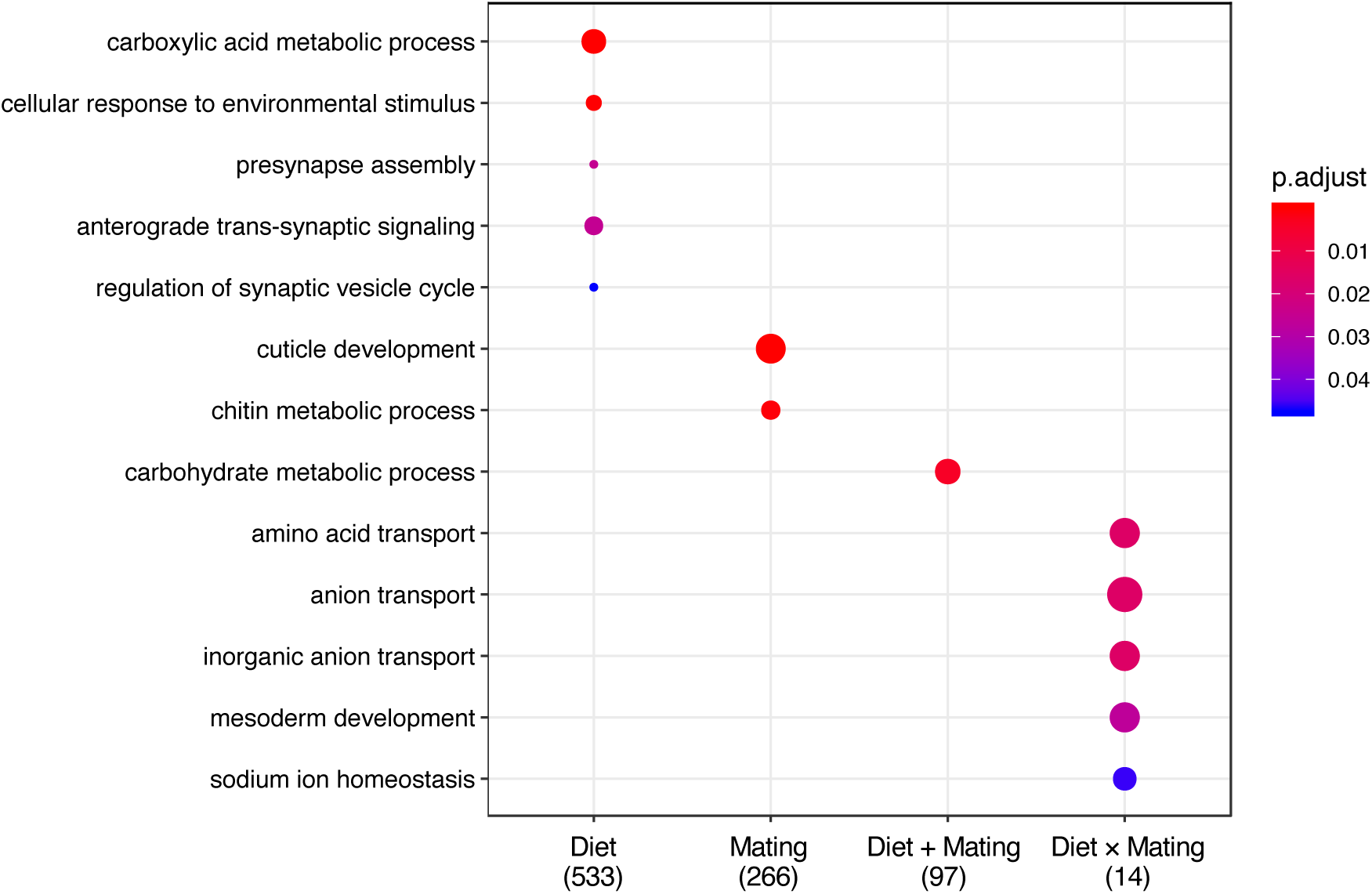
GO enrichment for the transcriptomic response to changes in diet and mating status. Enrichment analyses for ‘biological process’ were performed for all gene categories indicated on the x-axis, and p-values were adjusted for FDR < 0.05 (’p.adjust’).

**Table 2:** Gene overlap analysis between expression categories for female-expressed genes obtained from our current analysis (see Tab. 1) and categories defined in previous work investigating sex-specific expression responses to diet (Camus et al. 2019, ref. [4]).

|  |  | Current Female-specific Dataset |  |  |  |
| --- | --- | --- | --- | --- | --- |
|  |  | Diet | Mating | D + M | D × M |
|  |  | 651 | 345 | 110 | 14 |
| **Camus**<br>**et al.**<br>**(2019)** | Diet<br>639 | **305** | 19 | **47** | **7** |
|  | Diet × Sex<br>51 | **15** | 5 | **5** | 0 |
|  | D + (D × S)<br>116 | **70** | 2 | **31** | **5** |
|  | f. reproductive<br>165 | 8 | **45** | 0 | 0 |

Genes in the Mating set represent female responses to fertilisation. Fold-changes from virgin to mated state are mostly positive (indicating increased expression after mating) and—as expected—are highly correlated across diets (**Figure 1B**). The set of mating-dependent genes identified here overlapped significantly with those previously reported as showing differential expression between mated and virgin females in the abdomen ([20], overlap = 19 genes, p< 0.001), but not in head-thorax (overlap = 102 genes, p = 0.5). They are also significantly enriched for previously identified reproductive genes with female-limited expression ([4], overlap = 45 genes, p < 0.001, **Table 2**). Gene Ontology (GO) enrichment analysis revealed that mating-dependent genes enriched for terms related to cuticle development, chitin catabolic process and cell adhesion (**Figure 2**).

For the small Diet+Mating category we find significant overlap with genes previously shown to have concordant diet-responses across the sexes (D category in [4]: overlap = 47 genes, p< 0.001, **Table 2**), as well as sex-specific diet-dependent regulation (DxS category in [4]: overlap = 5 genes, p < 0.001; D+(D×S): overlap = 31 genes, p < 0.001, **Table 2**). The genes show concordant responses to mating between diets (r = 0.884, p < 0.001, **Figure 1C**) and concordant responses to diets between mating treatments (r = 0.811, p < 0.001, **Figure 1D**, possibly with some undetected carbohydrate-specific expression changes following mating). In terms of function, genes in the Diet+Mating category were significantly enriched for carbohydrate metabolic processes (**Figure 2**).

Finally, for the Diet×Mating interaction category, we find patterns of fold change that indicate genes that are up-regulated in the carbohydrate environment in mated flies only, while being unaffected by diet in virgins (**Figures 1E and 1F**). Despite its small size, we found significant overlap between the gene set and those previously shown to have concordant diet responses across the sexes (D category in [4]: overlap = 7 genes, p < 0.001, **Table 2**) and those showing opposing diet-dependent regulation between males and females (D×S category [4]: overlap = 5 genes, p < 0.001, **Table 2**). Functionally, Diet×Mating genes are significantly enriched for biological processes of amino acid transport (**Figure 2**).

### Expression regulation

We used analyses that detect the enrichment of binding motifs in the regulatory regions of sets of genes to infer the transcription factors that are the best candidates for driving the observed patterns of differential expression (**Table 3**, see data sets 6 and 7 in online repository [46] for full results).

**Table 3:**
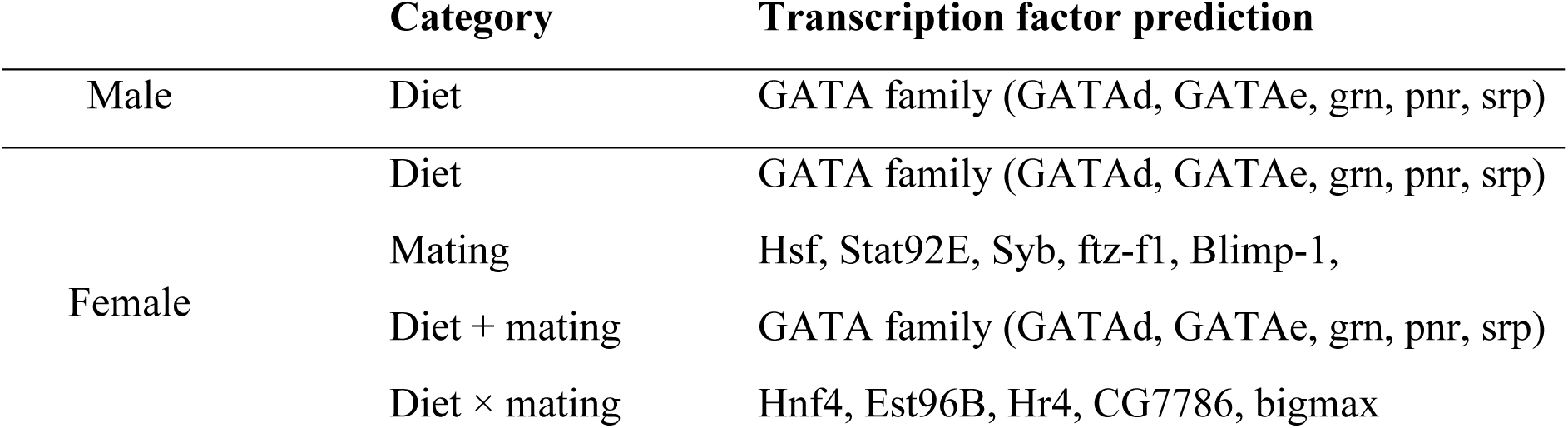
Enriched transcription factor binding motifs for each of the four expression categories defined in Table 1.

For males, the genes responding in expression to diet are enriched for transcription factors of the GATA family, such as *grain* (*grn*), *GATAd*, *GATAe*, *pointer* (*pnr*) and *serpent* (*srp*). In the more complex female dataset, we found both the Diet and the Diet+Mating sets to be enriched for regulation by transcription factors of the GATA family, similar to regulation in males. Genes in the Mating set were enriched in binding motifs for *Heat shock factor* (*Hsf*), as well as *Signal-transducer and activator of transcription protein at 92E* (*Stat92E*), *Synaptobrevin* (*Syb*), *ftz transcription factor 1* (*ftz-f1*), and *Blimp-1*. Finally, genes in the Diet×Mating category had upstream regions enriched for binding motifs for *Hepatocyte nuclear factor 4* (*Hnf4*), *Ets96B*, *Hormone receptor 4* (*Hr4*), CG7786, and *bigmax*.

### Thermal tolerance

With gene expression in response to mating enriched for regulation by *Hsf*, we assessed thermal tolerance in virgin and mated females on the two diets. The data showed that overall, mated females were significantly more heat-tolerant than virgin females (**Table SR1.4**, **Figure 3**). Additionally, we found that females fed on high-protein diet were also significantly more heat-tolerant than flies fed on high-carbohydrate diet (**Table SR1.4**, **Figure 3**). Both effects were of similar size (coefficients ± SE: mating = −2.86 ± 1.19, diet = −2.16 ± 1.17) and additive (mating status-by-diet interaction not significant, **Table SR1.4**). Accordingly, out of the four treatment groups, mated females fed with the protein-rich diet had the highest heat tolerance levels, with mated females on the carbohydrate-rich diet having a similar thermal tolerance to virgin flies on protein-rich diet and virgins fed the protein-rich diet having the lowest heat tolerance (**Figure 3**).

**Figure 3:**
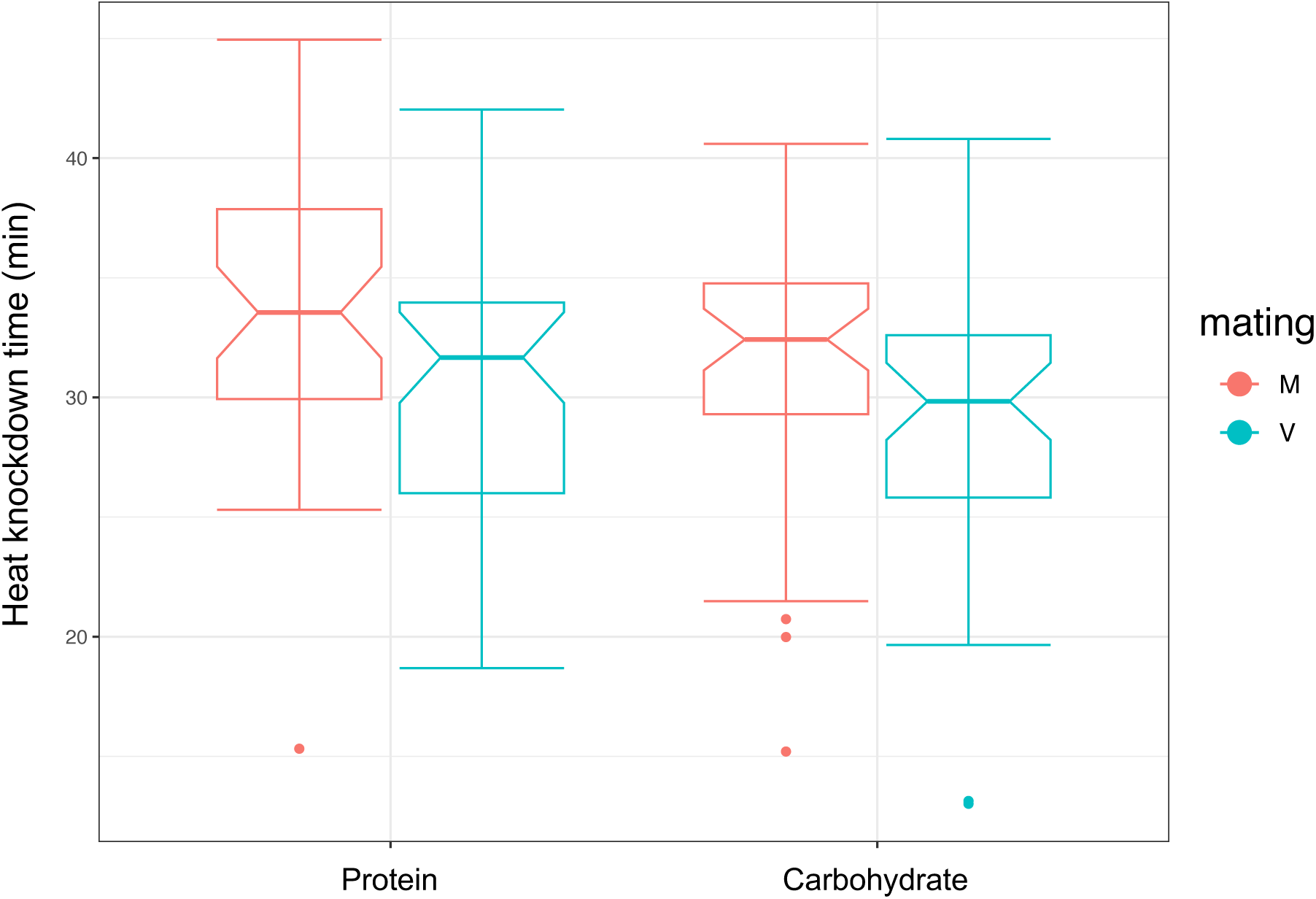
Boxplots of heat knockdown response (time to knock down, y-axis) of mated (red) and virgin (blue) females fed a protein-rich or a carbohydrate-rich diet (x-axis). Bold lines indicate medians, boxes the interquartile range, whiskers the most extreme values within 1.5 interquartile ranges. Dots indicate data points beyond this range.

## Discussion

In this study, we aimed to clarify the relationship between nutrition and reproduction in *Drosophila*. We placed virgin and mated flies into two nutritional environments that differed in their protein-to-carbohydrate ratios and had previously been shown to represent the female- and male-optimal diets [4]. We performed phenotypic measurements to confirm that our diet manipulations replicated previously described patterns and then investigated the transcriptome across these treatments to separate genes involved in nutritional responses from those that are regulated in line with reproductive investment.

Our phenotypic experiments replicated previous findings, thus validating our approach. In no-choice assays, we found the expected patterns of sex-specific fitness on the two types of diet across virgin and mated flies [5, 9, 16], where females are more fecund on protein-rich than carbohydrate-rich food while males gain greater mating success on the carbohydrate-rich food. Choice assays provided evidence for a limited response to mating of male diet preference [13, 24, 25], compared to a large shift of diet preference in females [13, 18].

Transcriptome analysis revealed sex differences in diet- and mating-dependent gene expression that match those observed in feeding behaviour. Males show a very restricted change in expression between the two types of diet, but this response is independent of mating status, and mating in itself does not result in any transcriptomic changes. Females, in contrast, not only share the core dietary response that is seen in males but also exhibit profound changes in gene expression patterns that are triggered by mating, as well as some mating-dependent changes in their responses to diet.

The fact that the male diet response includes many ejaculate-associated genes whose expression is regulated in parallel to reproductive output suggests that male reproductive investment is modulated in line with the nutritional environment. This contrasts with the absence of any plasticity in response to mating status, implying that adult males are in a reproductively active state whether mating takes place or not. This fact is consistent with an onset of spermatogenesis already in the pupal state [47]. It should be noted, however, that the absence of a transcriptional mating response in males reported here is at least somewhat at odds with the only two previous studies that have compared gene expression in virgin and mated males. Ellis and Carney [48] measured gene expression in heads using microarrays and found a modest response, with 47 genes differentially expressed between males that had mated vs. controls that had not been presented with a female. Fowler et al. [20] used RNA-seq to assay gene expression in two tissues, head-thorax and abdomen. While they detected no expression change in head-thorax, they found a much larger response to mating in the abdomen with 2068 differentially expressed genes [20]. The fact that these effects of mating were not detected in our study could be due to signals being obscured in the whole flies (rather than specific tissues) analysed here. Furthermore, differences could arise due to the timing of samples collection.

The two other studies were conducted by taking samples soon after an observed mating (2 hours for Ellis and Carney, 3h for Fowler et al.). It is possible that expression changes induced by mating are transient and could then have been missed in our design, where males were sampled from mating groups independently of the occurrence of recent mating events. It is worth noting, however, that the differential expression of accessory and seminal fluid proteins between mated and virgin males that was detected by Fowler et al. ([20], table S2d), presumably due to replenishment after mating, is also apparent in our dataset, but as correlate to differences in overall reproductive investment between the diet treatments.

In contrast to results in males, the large transcriptional response to mating that we observe in females is in line with previous results. For example, Fowler et al. [20] found 125 genes to be differentially expressed in response to mating in the female abdomen and 2040 genes in head-thorax. Similarly, Delbare et al. [49] compared the mating response across lines of different geographic origins and found a shared set of 272 genes that were consistently differentially expressed between mated and virgin females (in addition to a further 77 genes that showed differential expression in specific combinations of female and male origin). Beyond these quantitative similarities, we also find correspondence between the specific genes that we and previous work identify as being regulated in response to mating. Thus, genes in our Mating category overlap significantly with those that Fowler et al. [20] detected as differentially expressed between the abdomens of mated and virgin females. We note that a similar result was not obtained when comparing our gene set with Fowler et al.’s differentially expressed genes from head-thorax. However, it is worth noting that their gene set is very large (more than 2000 genes) and contains a smaller proportion of genes that are also detected as mating-responsive in other studies than the thorax set (29% vs. 65% of genes also detected elsewhere [20]).

While expression changes associated with mating have been well studied, we were also able to assess the effect of diet alongside and in combination with the mating response. Doing so, we were able to generate several new insights. First, our analysis allowed us to extend the inference of a core metabolic response to macronutrient composition from mated females and males (4) to virgin females and males. The existence of such a fundamental physiological response to diet is not surprising, but the fact that it also occurs in reproductively inactive individuals corroborates this finding. More interestingly, our analysis of transcription factor binding sites revealed that the core response to diet composition, shared between virgin and mated females as well as males, is heavily reliant on transcription factors of the GATA family. This could seem surprising, as members of this transcription factor family have been associated with a number of reproductive and ageing phenotypes. Thus, GATA transcription factors are involved in the regulation of reproductive genes in fruit flies [50] and mosquitoes [51], as well as in the ageing response to dietary restriction in *Caenorhabditis elegans* [52] and regulation associated with dietary restriction and rapamycin treatment in flies [7]. While these functions should not apply to virgins, our results are more in line with previous studies that implicated GATA factors in the regulation of physiology and feeding, especially in response to sugar [53]. Further work will be needed to investigate whether the regulation of life history decisions and more general physiology are distinct functions of GATA factors, or whether regulators of physiology are deployed in different contexts.

Beyond the core response to diet composition, the analyses presented here suggest that diet composition and mating responses act largely independently and additively, with the number of genes for which we detect a significant diet-by-mating status interaction being very small. This is not what we had expected, as we had predicted that diet and mating would act synergistically. Specifically, we had expected that a significant number of genes would show differential responses to diet in virgin and mated females, with a larger increase in expression under the more favourable conditions in mated females than in virgins, matching feeding rates and reproductive investment. The fact that we observe an additive effect of mating would imply that the mating response is largely qualitative (mating induces the production of gene products) rather than quantitative (the amount of gene product produced scales with reproductive output).

When interpreting the above results, however, it is important to keep in mind that the number of genes with a significant interaction effect between diet and mating status is likely to be limited by the statistical power to pick up these more subtle effects. It is interesting in this respect to contrast the findings of our factorial analysis with results obtained by separate univariate analyses performed in each mating regime (see **Supplementary Results 2**). These analyses detect a large excess of differentially expressed genes in the mated compared to the virgin mating regime (887 vs. 102 genes). Of course, this result is also limited by statistical power, which here reduces the ability to detect shared diet-specific regulation when analysing two datasets of half the size compared to the joint analysis in the main text. So taken together, these results suggest that there is a sizeable set of genes that show quantitative differences between mated and virgin flies in their level of diet-specific regulation, where fold changes are larger in the former and smaller in the latter. These differences in differential expression are small enough to not result in significant interaction effects in the joint analysis, but large enough to lead to contrasting significance in the separate analyses.

Another interesting result from this work is the identification of *Heat shock factor* (*Hsf*) as enriched in the regulation of genes related to reproduction and diet. As their name suggests, heat-shock factors were first described as drivers for responses to thermal stress [54]. However, there have been several studies suggesting roles in important biological functions in the absence of stress. Most notably, *Hsf* is required for oogenesis in *Drosophila* in a way that does not depend on the expression of heat-shock proteins [55] and shows its highest level of tissue-specific expression in ovaries [56]. A similar role for heat-shock factors in fecundity has been described in mice [57–59], where they also have been shown to play a role in sperm production (50). In line with the role of heat-shock factors in reproduction, a large number of *Hsf* binding sites in *Drosophila* are associated with genes that are not involved in the heat-shock response and *Hsf*-regulated genes are enriched for GO categories associated with gamete formation and oogenesis [60]. In light of these previous findings, it seems that *Hsf* is involved in the diet-dependent regulation of female reproduction, binding to genes that are differentially regulated in response to the macronutrient composition of the food consumed by females.

Not much is known about the mechanistic links between *Hsf* and diet. Work in rodent models has also shown that the amino acid glutamine is involved in the activation of *Hsf* [61], while experiments with the nematode *C. elegans* have demonstrated links between *Hsf* and the insulin-signalling network, with inputs from both being required for development, stress response, survival and reproduction [62]. In *Drosophila*, larvae raised in high-protein environments are better able to tolerate thermal stress than flies raised in carbohydrate-rich environments [63, 64]. A recent study further found that males fed a control food take longer to be knocked down by heat than males fed a restricted diet rich in complex carbohydrate– cellulose [65]. However, no study to date has examined how mating status interacts with the dietary environment to shape the heat-shock response. Our experiments confirmed a role of diet in modulating heat resistance, showing that flies fed a protein-rich diet were more heat tolerant than flies fed a carbohydrate-rich food, possibly because a protein-rich diet favours the production of protective proteins such as heat-shock proteins, antioxidants, and enzymes involved in repair. Moreover, we found that independently of the diet treatment, mated flies exhibited better tolerance to heat compared to virgins, suggesting that the metabolic conditioning induced post-mating extends beyond egg or offspring production, generating additional protective mechanisms without the trade-offs previously assumed. Future studies should dissect this further by editing one or more known heat-shock factors to determine which are specifically involved in oogenesis versus those contributing to stress tolerance. It is unclear whether and to which degree these two roles are related. Previous work has proposed that the role of *Hsf* in oogenesis is independent of the heat-shock response [55], yet our observations suggest some possible links.

In conclusion, our aim was to investigate the metabolic pressures placed upon an organism as a consequence of reproduction. We find that in female *Drosophila*, there is a large metabolic rewiring that occurs after mating, and this is mainly to fuel the process of oogenesis. We were able to dissect the nutritional response into basal and reproduction-related categories and identified several transcription factors as possible drivers of these responses, including GATA family transcription factors and *Hsf*. Our work provides further understanding on how organisms are able to cope with changes in the internal and external physiology.

## Supporting information

Supplementary Results 1

Supplementary Results 2

## Acknowledgements

We would like to thank Nazif Alic and Adam Dobson for their help with planning the experiments, Matt Piper for his input into the interpretation and Rebecca Finlay for help with laboratory work.

