## Supplementary Results 1 for "Disentangling the reproductive and metabolic transcriptional responses to diet in *Drosophila melanogaster*"

### Supplementary Results 1: Phenotypic analyses

**Figure SR1.1: Phenotypic responses to nutrition. (A)** Consumption of liquid protein and carbohydrate diets (as mean intake ± standard error) in diet preference assays. Data are shown for virgin and mated flies of both sexes, with symbols and colours as described in the legend. The dashed line represents the 1:1 ratio **(B)** Total liquid diet consumption in no-choice assays. Females are shown in the left panel and males on the right, with mating status depicted by the two colours as shown in the legend of panel A. **(C)** Fitness values for females (competitive fecundity – left) and males (competitive fertilisation success – right) across both nutritional environments. Mating status is shown by the two different colours as shown in the legend of panel A.

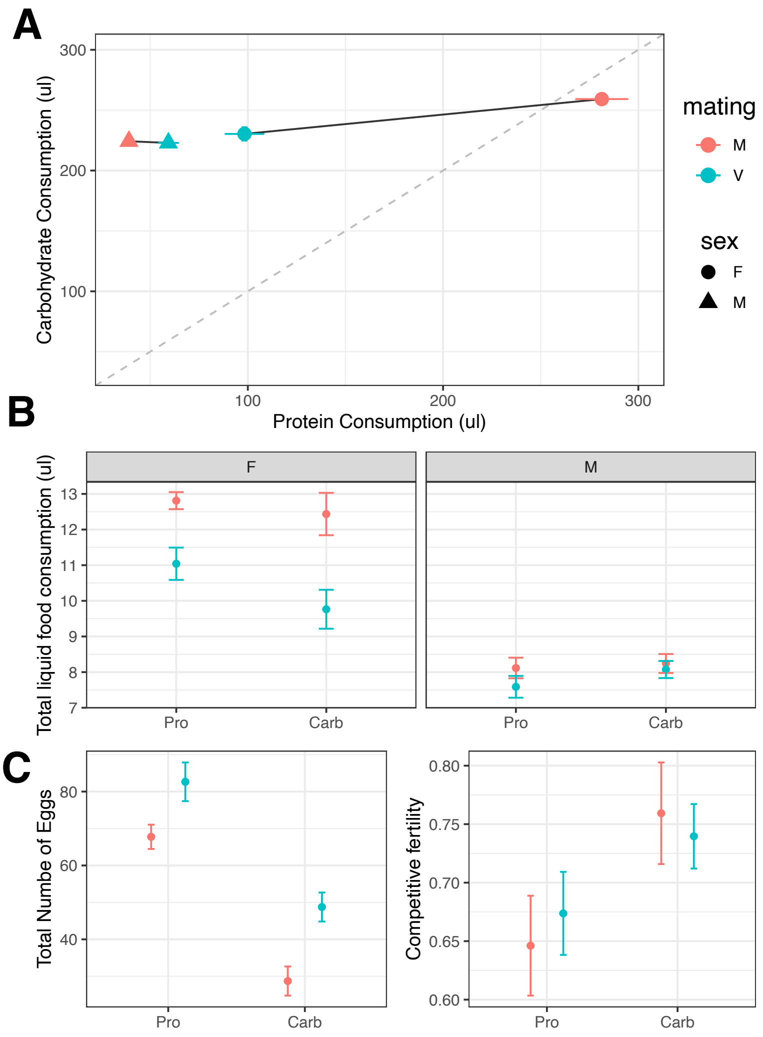

**Table SR1.1: Consumption of protein and carbohydrate food in diet preference assays.** Results from multivariate (MANOVA) and univariate (ANOVA) analyses of the total amount of liquid (µl) consumed by triplets of flies. Predictor variables are sex, mating status (virgin/mated) and their interaction.

**Overall Model**

| | Df | Pillai | approx F | numDf | denDf | Pr(>F) |
|---|---|---|---|---|---|---|
| sex | 1 | 0.79285 | 143.526 | 2 | 75 | < 0.001 |
| mating | 1 | 0.5794 | 51.659 | 2 | 75 | < 0.001 |
| sex × mating | 1 | 0.6601 | 72.826 | 2 | 75 | < 0.001 |
| Residuals | 76 | | | | | |

**Response protein:**

| | Df | Sum Sq | Mean Sq | F value | Pr(>F) |
|---|---|---|---|---|---|
| sex | 1 | 395129 | 395129 | 228.576 | < 0.001 |
| mating | 1 | 132506 | 132506 | 76.652 | < 0.001 |
| sex × mating | 1 | 206483 | 206483 | 119.447 | < 0.001 |
| Residuals | 76 | 131378 | 1729 | | |

**Response carbohydrate:**

| | Df | Sum Sq | Mean Sq | F value | Pr(>F) |
|---|---|---|---|---|---|
| sex | 1 | 8980.6 | 8980.6 | 28.578 | < 0.001 |
| mating | 1 | 4563.8 | 4563.8 | 14.523 | < 0.001 |
| sex × mating | 1 | 3781.9 | 3781.9 | 12.035 | < 0.001 |
| Residuals | 76 | 23882.7 | 314.2 | | |

**Table SR1.2: Analysis of diet consumption from no choice dietary assays.** The table shows the Analysis of Variance of a linear models with total liquid food consumed as a response variable and sex, mating status, diet treatment and all their interactions as fixed effects.

| | Sum Sq | Df | F value | Pr(>F) |
|---|---|---|---|---|
| sex | 413.75 | 1 | 114.5471 | < 0.001 |
| mating | 38.62 | 1 | 10.6931 | 0.001334 |
| diet | 0.01 | 1 | 0.0037 | 0.951354 |
| sex × mating | 16.87 | 1 | 4.6716 | 0.032252 |
| sex × diet | 3.34 | 1 | 0.9258 | 0.337493 |
| mating × diet | 6.41 | 1 | 1.7739 | 0.184921 |
| sex × mating × diet | 13.75 | 1 | 3.8055 | 0.052947 |
| Residuals | 541.81 | 150 | | |

**Table SR1.3: Analyses of female and male reproductive fitness in no choice dietary assays.** The table shows the Analysis of Variance of linear models with sex-specific reproductive fitness as a function of total food consumption (covariate, see Table SR1.2), diet, mating status and their interaction as fixed effects. Female fitness was measured as total number of eggs laid in an 18-hour period, male fitness as the proportion of offspring sired in competition with a standard male.

**(a) Female fitness**

| | Df | Sum Sq | Mean Sq | F value | Pr(>F) |
|---|---|---|---|---|---|
| diet | 1 | 25907.0 | 25907.0 | 75.4177 | < 0.001 |
| status | 1 | 5090.9 | 5090.9 | 14.8202 | < 0.001 |
| total consumption | 1 | 2.7 | 2.7 | 0.008 | 0.929031 |
| diet × status | 1 | 115.6 | 115.6 | 0.3365 | 0.5636268 |
| Residuals | 73 | 25076.4 | 343.51 | | |

**(b) Male fitness**

| | Df | Sum Sq | Mean Sq | F value | Pr(>F) |
|---|---|---|---|---|---|
| diet | 1 | 109.2 | 109.2 | 6.2229 | 0.01481 |
| status | 1 | 0.5 | 0.5 | 0.029 | 0.86519 |
| total consumption | 1 | 7.5 | 7.5 | 0.4295 | 0.51423 |
| diet × status | 1 | 12.5 | 12.5 | 0.7133 | 0.40104 |
| Residuals | 75 | 1316.2 | 17.6 | | |

**Table SR1.4: Analyses of female thermal resistance.** The table shows the Analysis of Variance for the fixed effects of a linear mixed model of time until heat knockdown as a function of mating status ('status'), diet and their interaction.

| | Sum Sq | Mean Sq | NumDF | DenDF | F value | Pr(>F) |
|---|---|---|---|---|---|---|
| status | 279.248 | 279.248 | 1 | 169.91 | 9.4091 | 0.002513 |
| diet | 146.308 | 146.308 | 1 | 169.06 | 4.9297 | 0.027728 |
| status × diet | 5.255 | 5.255 | 1 | 170.12 | 0.1771 | 0.674429 |
