## Supplementary Results 2 for "Disentangling the reproductive and metabolic transcriptional responses to diet in *Drosophila melanogaster*"

### Supplementary Results 2: Additional RNA-seq analyses

We performed an alternative analysis of the transcriptomic data for virgin and mated females, to identify genes that are part of the core response to diet consumption versus those that are related to mating status. Rather than running a model with diet, mating status and their interaction on the entire dataset, we separately analysed the data for virgins and mated females, testing for an effect of diet (**Figure SR2.1A**). This approach assumes that the nutritional response in virgin flies provides the baseline for the physiological response to diet, whereas mated flies show the baseline response to diet plus the additional transcriptomic signature of reproduction.

We found 102 genes to respond to dietary change in virgin flies, whereas mated flies had 887 differentially expressed genes (**Figure SR2.1B**). In line with our prediction, we found a large overlap between virgin and mated nutritional responses, with 86 genes being shared, encompassing most of the virgin response (**Figure SR2.1B**). Among these genes, carbohydrate-to-protein fold changes were strongly positive correlated between virgins and mated females ($r = 0.835$, $p < 0.001$), indicating that the basal response to nutrition is not dependant on mating status.

Given the large overlap between the virgin and mated datasets, we subdivided the data into two main categories (**Figure SR2.1B**), the basal diet response (the 102 genes with differential expression between diets in virgin flies) and the set of genes that respond to diet only in mated flies (comprised of the 801 genes that showed differential expression in mated flies, excluding the 86 genes overlapping with the basal response). The basal and the mated diet responses show significant differences in the direction of gene regulation ($Chi^2 = 67.73$, df = 1, $p < 0.0001$), where basal genes are mostly upregulated on protein-rich relative to carbohydrate-rich food, while genes specifically regulated in mated flies are in majority down-regulated (**Table SR2.1**).

Analyses of overlap between gene sets defined in the analyses presented here and those obtained by the global analysis of female gene expression in the main text (**Table SR2.2**) show that the two analyses split differentially expressed genes differently. All categories defines here (Basal and Mated, or Basal, Mated-up and Mated-down) overlap with both the Diet and the Diet+Mated categories defined in the main text, with the larger Mated and Mated-down sets also overlapping with the Diet×Mating set. Thus, some genes that only show differential expression in response to diet in mated flies in the analyses here were detected as showing a general diet response across all individuals in the global analysis, rather than a diet-by-mating interaction. As discussed in the main text, these differences most likely arise due to different limitations in power between the two analyses.

GO analyses show that the genes that constitute the basal diet response are highly enriched for several metabolic processes (**Figure SR2.2**), while analyses of transcription factor binding site enrichment indicates regulation by GATA family transcription factors (**Table SR2.2**), just like genes of the male and female Diet categories in the analyses of the main text (**Table 3**).

**Figure SR2.1: Alternative analysis of female diet and mating responses. (A)** Experimental design of contrasts used. In brief, we examined nutritional changes (going from carbohydrate to protein) in virgin and mated female separately. **(B)** Gene overlap for genes that respond to nutritional changes in virgin and mated flies.

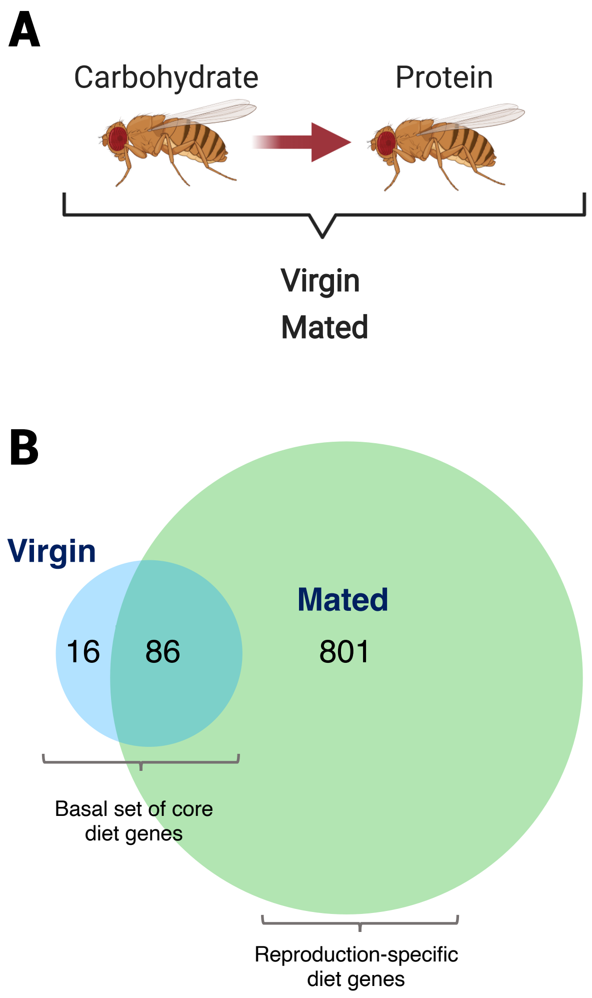

**Figure SR2.2:** GO enrichment for the transcriptomic response to changes in nutritional environment and mating status. Enrichment for 'biological process' was performed for all categories, and p-values were adjusted for FDR < 0.05 ('p.adjust').

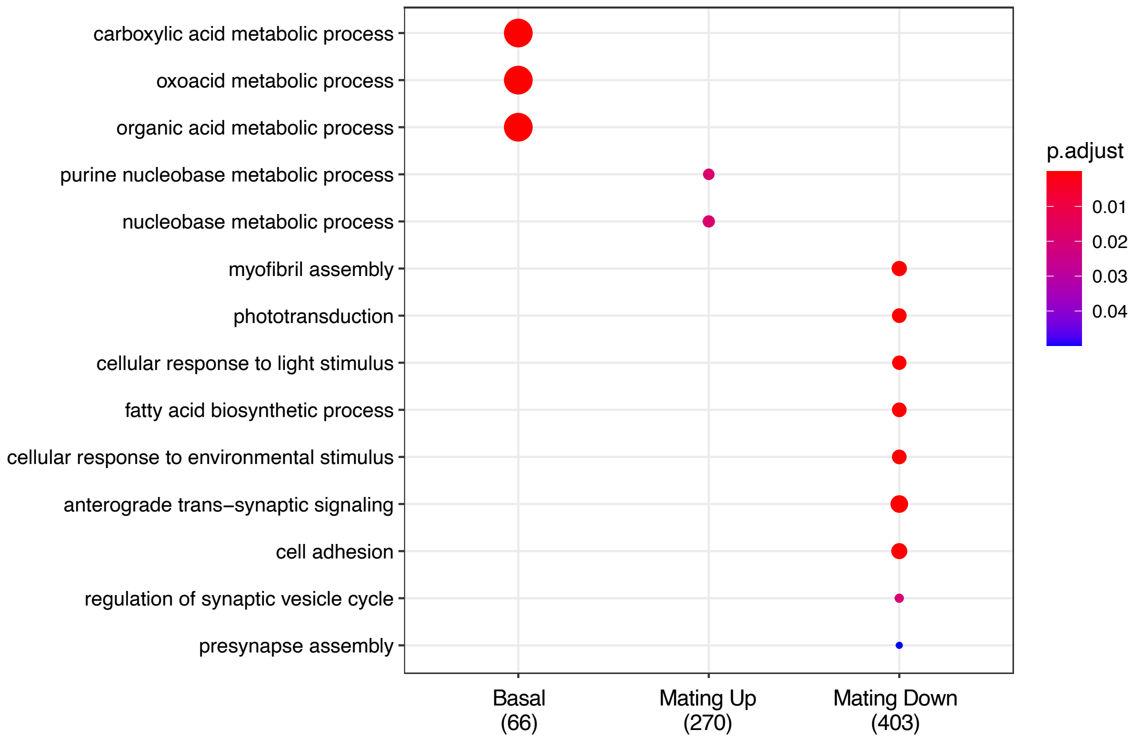

**Table SR2.1:** Number of genes with significant up- or down-regulation between a carbohydrate-rich and a protein-rich diet in virgin and mated flies. Expression changes in virgin flies is designated as the basal response to diet.

| | Up | Down |
|---|---|---|
| Basal (virgin) | 84 | 19 |
| Mated | 307 | 494 |

**Table SR2.2:** Gene overlap analysis between categories of female gene regulation obtained from the supplementary analyses of virgin and mated flies presented here (columns) and those obtained in the global analysis presented in the main text (rows). The number of genes in each category is given in the column and row labels, the number of genes that overlap between pairs of categories is shown in the table cells. Significant overlap is indicated by bold face.

| | Basal (102) | Mated (801) | Mated up (307) | Mated down (494) |
|---|---|---|---|---|
| Diet (651) | **66** | **435** | **158** | **277** |
| Mating (345) | 0 | 22 | 8 | 14 |
| D+M (110) | **35** | **57** | **35** | **22** |
| DxM (14) | 1 | **13** | 0 | **13** |

**Table SR2.3:** Enriched transcription factor binding motifs for each of the three categories.

| **Category** | **Transcription factor prediction** |
|---|---|
| Basal | GATA family (GATAd, GATAe, grn, pnr, srp) |
| Mated up | Hsf, pb, GATA family (grn, srp) |
| Mated down | Optix, NfI, Mef2 |
