## Supplementary Methods for "Disentangling the reproductive and metabolic transcriptional responses to diet in *Drosophila melanogaster*"

### Supplementary Methods: Composition of diets

**Table SM.1:** Composition of the standard cornmeal-agar-molasses diet. The recipe is given for 3l of media.

| Ingredient | Quantity |
|---|---|
| Molasses | 200ml |
| Agar | 24g |
| Cornmeal | 200g |
| Yeast powder | 82g |
| Nipagin (100mg/L) | 90ml |
| Propionic Acid | 9ml |

**Table SM.2:** Essential and non-essential amino acid stock solutions.

| Amino acid stock solution | **(g/200 ml)** |
|---|---|
| **Essential amino acid** | |
| F (L-phenylalanine) | **3.03** |
| H (L-histidine | **2.24** |
| K (L-lysine) | **5.74** |
| M (L-methionine) | **1.12** |
| R (L-arginine) | **4.70** |
| T (L-threonine) | **4.28** |
| V (L-valine) | **4.42** |
| W (L-tryptophan) | **1.45** |
| **Non-essential amino acid** | |
| A (L-alanine) | **5.25** |
| D (L-aspartate) | **2.78** |
| G (glycine) | **3.58** |
| N (L-asparagine) | **2.78** |
| P (L-proline) | **1.86** |
| Q (L-glutamine) | **6.02** |
| S (L-serine) | **2.51** |

**Table SM.3:** Recipe for 200ml of protein solution

| | | | **Total volume 200ml** |
|---|---|---|---|
| | L-ile | Powder | 348mg |
| | L-leu | Powder | 492mg |
| | L-tyr | Powder | 252mg |
| | cholesterol | 20mg/ml in EtOH | 3ml |
| | CaCl2 | 1000x | 200ul |
| | MgSO4 | 1000x | 200ul |
| | CuSO4 | 1000x | 200ul |
| | FeSO4 | 1000x | 200ul |
| | MnCl2 | 1000x | 200ul |
| | ZnSO4 | 1000x | 200ul |
| | $H_2O$ | | Up to 50ml |
| Total volume before autoclaving | | | 50 ml |
| | buffer | 10x acetate buffer base | 20ml |
| | nucl/lipid soln | 125x stock | 1.6ml |
| | Yaa solutions | essential amino acid stock solution (EAA) | 18.154ml |
| | | non essential amino acid stock solution (NEAA) | 18.154ml |
| | | Na glutamate solution (100mg/ml) | 5.464ml |
| | | Cys solution (50mg/ml) | 1.584ml |
| | Vitamin stock | 47.6x stock | 4.2ml |
| | folic acid stock | 1000x stock | 200ul |
| | Propionic acid | | 1.2ml |
| | Nipagin | 100 g/l stock in 95% EtOH | 3ml |
| Make to total volume of 200ml with $H_2O$ | | | |

**Table SM.4:** Recipe for 200ml of carbohydrate solution

| | | | Total volume 200ml |
|---|---|---|---|
| | sucrose | To match protein 1:1 | 6.5g |
| | cholesterol | 20mg/ml in EtOH | 3ml |
| | CaCl2 | 1000x | 200ul |
| | MgSO4 | 1000x | 200ul |
| | CuSO4 | 1000x | 200ul |
| | FeSO4 | 1000x | 200ul |
| | MnCl2 | 1000x | 200ul |
| | ZnSO4 | 1000x | 200ul |
| | $H_2O$ | | Up to 50ml |
| Total volume before autoclaving | | | 50ml |
| | buffer | 10x acetate buffer base | 20ml |
| | nucl/lipid soln | 125x stock | 1.6ml |
| | Vitamin stock | 47.6x stock | 4.2ml |
| | folic acid stock | 1000x stock | 200ul |
| | Propionic acid | | 1.2ml |
| | Nipagin | 100 g/l stock in 95% EtOH | 3ml |
| Make to total volume of 200ml with $H_2O$ | | | |
